# Genetic characterisation and aggressiveness of the *Fusarium oxysporum* Species Complex in tomato plants and irrigation water from Australian processing fields

**DOI:** 10.64898/2026.05.24.727468

**Authors:** Hanyue Feng, Sophia E. Callaghan, Jiajing Cao, Paul W. J. Taylor, Sigfredo Fuentes, Alexis Pang, Yu Pei Tan, Niloofar Vaghefi

**Affiliations:** School of Agriculture, Food and Ecosystem Sciences, Faculty of Science, the University of Melbourne, Parkville, Victoria 3010, Australia; Department of Agriculture, Fisheries and Forestry, Mascot, NSW 2020, Australia; Centre for Crop Health, University of Southern Queensland, Toowoomba, Queensland 4350, Australia; Queensland Department of Primary Industries, Dutton Park, Queensland 4102, Australia

**Author notes:** **Correspondence**: Niloofar Vaghefi.

**Keywords:** soilborne pathogens, yield decline, phylogeny, epidemiology, genetic diversity, pathogenicity

## Abstract

*Fusarium oxysporum* Species Complex (FOSC) includes important soilborne pathogens of a range of crops worldwide. In Victoria (VIC) and New South Wales (NSW), Australia, FOSC has been identified as the cause of stunting and poor growth in processing tomato fields, resulting in significant yield losses. A total of 40 FOSC isolates were obtained from symptomatic tomato plants, irrigation water samples and culture collections, which were confirmed to be FOSC by multi-gene phylogenetic analyses based on four genomic loci: beta tubulin, calmodulin, the second largest subunit of nuclear RNA polymerase II, and translation elongation factor one-alpha. There was a high level of genetic diversity among isolates, with multiple phylogenetic lineages detected. Glasshouse bioassays demonstrated that all isolates were pathogenic to processing tomato, resulting in significant reductions in plant growth, with above ground height reduced by 11 to 26% and root dry weight by 44 to 83% compared with the control (p < 0.05). Although aggressiveness varied among isolates, growth reduction occurred irrespective of their phylogenetic placement. Moreover, the shared genetic background of isolates from irrigation water and plant samples highlights the role of irrigation water as a potential source of inoculum in *Fusarium* epidemics, underscoring its significance for disease management in Australian processing tomato systems.

## 1. Introduction

Tomato is one of the most important vegetable crops globally, grown in variable climatic conditions and soil profiles (Zhu & Miller, 2025). There are two major types of cultivated tomatoes: one for fresh consumption, for example salad tomato that is usually grown under controlled environments (Cammarano *et al*., 2022); and processing tomato used for canned products, ketchup and sauces that is mostly grown under field conditions (Ma *et al*., 2023). The major production areas of processing tomatoes include California, the USA (Swett *et al*., 2023), Italy (Caruso *et al*., 2022), and China (Yang *et al*., 2023).

In Australia, tomato is the second-most profitable vegetable commodity (McKenzie *et al*., 2017). In 2022-2023 growing season, 321,736 tonnes of tomatoes were produced, valued at $570.6 million with 34% designated for the processing industry (Stewart, 2023). Approximately 2,000 to 3,000 hectares of processing tomatoes are produced annually, of which 75% is in northern Victoria (VIC) and the remaining 25% in New South Wales (NSW) (Ma *et al*., 2023). Over the past decade, an estimated 10% annual yield loss in Australian processing tomato production has been attributed to soilborne diseases (Callaghan *et al*., 2018), with *Fusarium oxysporum* identified as a major contributing factor. Members of the *Fusarium oxysporum* species complex (FOSC) are among the most destructive soilborne pathogens affecting tomato crops worldwide (Swett *et al*., 2023, Ramsey *et al*., 1992). While *F. oxysporum* is well known for its persistence in soil, recent studies suggest that irrigation water may also serve as a potential source of FOSC inoculum, particularly with the increasing use of recycled or surface water in agricultural systems (McGovern, 2015). This highlights the importance of considering both soil- and waterborne pathways in the epidemiology and management of *F. oxysporum* in processing tomato production.

In recent years, extensive research on FOSC has led to recognition of numerous cryptic species based on phylogenetic species concept, which ranges from three to fifteen species (Achari *et al*., 2020, O’Donnell *et al*., 1998, Lombard *et al*., 2019, Laurence *et al*., 2012). The molecular taxonomy of *F. oxysporum* has consequently undergone several revisions, with multiple frameworks proposed by different research groups. These efforts have been further complicated by evidence of horizontal gene transfer and genetic recombination within the complex (Zhang *et al*., 2020). Within FOSC, while isolates are traditionally grouped into *formae speciales* (f. sp.) based on host specificity (Lievens *et al*., 2008), these phenotypic classifications cannot be reliably inferred from phylogeny or effector gene profiles alone, as the definition of *forma specialis* (f. sp.) is phenotypic, relying on the host range and differential pathogenicity assays rather than genetic relatedness (Baayen *et al*., 2000). Genetically similar isolates can therefore differ in pathogenicity and host interactions, and isolates assigned to the same *f. sp.* may fall into distinct phylogenetic clades (Afordoanyi *et al*., 2022). This highlights that understanding FOSC requires integrating both genetic and phenotypic analyses, with experimental assessment of pathogenicity remaining essential to characterize isolate diversity and impact.

Historically, O’Donnell *et al*. (1998) used translation elongation factor 1-alpha (*tef1-α*) and the mitochondrial small subunit of the ribosomal RNA genes (*mtSSU rDNA*) to test the monophyly of several *F. oxysporum* isolates. Their analyses revealed that although *F. oxysporum* formed a monophyletic clade, both pathogenic and non-pathogenic *F. oxysporum* isolates occurred within the same evolutionary lineages. Moreover, many *formae speciales*, including f. sp. *cubense* causing Panama disease on banana, and f. sp. *lycopersici* and *radicis-lycopersici* associated with tomato diseases, were polyphyletic, with several isolates having independent evolutionary origins. This suggested that host pathogenicity evolved convergently in *F. oxysporum*. Similarly, a later study by Laurence *et al*. (2012) used the same genes: *tef-1α* and *mtSSU rDNA* to genetically characterise Australian *F. oxysporum* isolates collected from native ecosystems representing four different climatic conditions using the same dataset as proposed by O’Donnell *et al*. (1998). The Australian isolates were separated into five clades that indicated high genetic diversity (Laurence *et al*., 2012), however, host pathogenicity of the isolates were not assessed.

More recently, Lombard *et al*. (2019) used four genes: beta tubulin (*tub2*), calmodulin (*cmdA*), the second largest subunit of nuclear RNA polymerase II (*rpb2*), *and tef-1α* for the multi-locus phylogenetic inference and combined with morphological studies to resolve cryptic speciation within FOSC and described fifteen separate species within eight major clades. Later on, Achari *et al*. (2020) applied multi-species coalescent analyses to whole genomes, nuclear genes, and mitochondrial genome sequence data to reconstruct the phylogeny of *F. oxysporum* isolates from both natural and agricultural ecosystems in Australia. Three major phylogenetic clades were consistently identified across all phylogenies, with two additional single-isolate clades. Therefore, it was suggested that the three clades may be considered as separate species, with Clade 3 representing *F. oxysporum sensu stricto* as it contained the neotype of *F. oxysporum*.

*Secreted In Xylem* (*SIX*) genes have been widely studied as key effectors contributing to virulence and host-pathogen interactions within the *Fusarium oxysporum* Species Complex, particularly in well-characterised pathosystems such as *F. oxysporum* f. sp. *lycopersici* (Jangir *et al.,* 2021). These genes, often located on lineage-specific or mobile pathogenicity chromosomes, can play important roles in determining compatibility with specific hosts and in triggering host resistance responses. However, increasing evidence indicates that the presence, absence, or diversity of *SIX* genes does not consistently align with phylogenetic relationships or reliably delineate *formae speciales* across the FOSC (Afordoanyi *et al*., 2022). Recent work has highlighted that host specificity in *F. oxysporum* may be more flexible and context-dependent than previously assumed, with multiple genetic lineages capable of infecting the same host and individual *formae speciales* not representing discrete evolutionary units (Han *et al*., 2025). As a result, while *SIX* genes provide valuable insight into pathogenic mechanisms, they are not currently incorporated into taxonomic frameworks for FOSC. Accordingly, the present study focuses on multi-locus phylogenetic analyses and pathogenicity assays to resolve taxonomic relationships among *F. oxysporum* isolates associated with processing tomato in Australia, rather than on effector profiling.

The phylogenetic placement of Australian *F. oxysporum* isolates recovered from processing tomato plants, along with their pathogenicity and aggressiveness toward local cultivars, remains poorly understood. To address this knowledge gap, the present study aimed to: 1) recover and genetically characterise *F. oxysporum* isolates associated with poor growth of processing tomatoes from affected fields in VIC and NSW, Australia; 2) investigate their phylogenetic relationships using different taxonomic frameworks; and 3) evaluate the pathogenicity and aggressiveness of representative isolates through replicated glasshouse bioassays. In this context, pathogenicity refers to the ability of an isolate to cause disease, whereas aggressiveness denotes the quantitative degree of disease severity expressed by a pathogenic isolate (De Silva *et al*., 2021).

## 2. Materials and methods

### 2.1. Sample collection and pathogen isolation

During the 2022, 2023, and 2024 growing seasons, a total of 28 symptomatic processing tomato plants were collected from 20 commercial fields in VIC and NSW, along with six irrigation water samples from one field in NSW. For plant sampling, whole tomato plants showing stunting, chlorosis, and poor growth (Figure 1) were carefully uprooted and transported to the laboratory. Samples were stored at 5 °C for a maximum of 48 hours before fungal isolation. For irrigation water sampling, six samples along five points of the irrigation system were collected; these included point 1: one sample from irrigation channel water as it entered the property; point 2: one sample from on farm reservoir filled with water from irrigation channel; point 3: one sample from on farm reservoir filled with rain and irrigation runoff; point 4: one sample from filtered water after leaving point 2 and before entering the irrigation system; and point 5: two water samples from irrigation drip lines in the field (Table 1).

**Figure 1.**
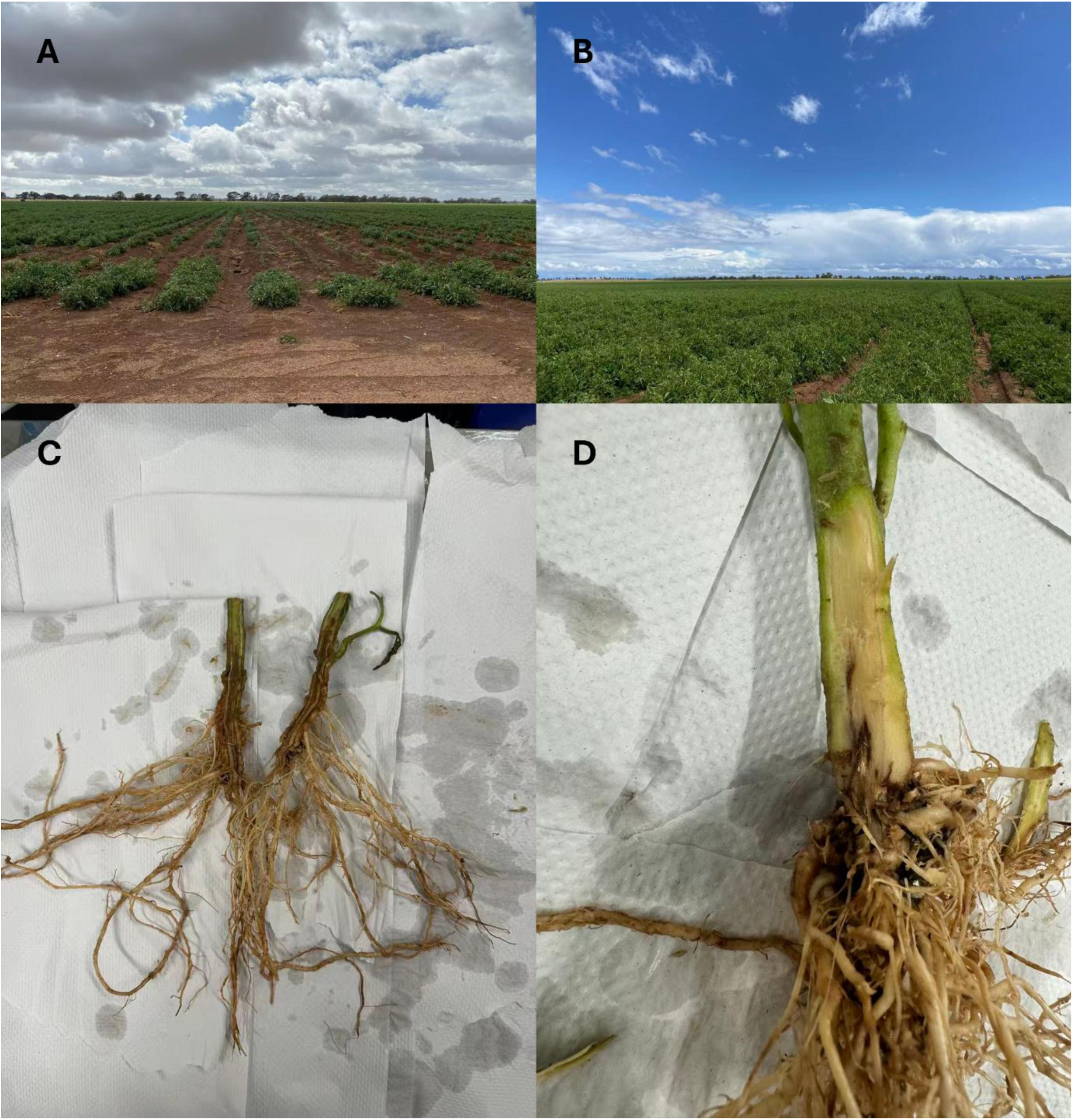
Comparison of (A) a disease-affected field with severe patchiness with a (B) healthy processing tomato field. (C and D) Collar, root, and shoot regions of *Fusarium oxysporum* infected processing tomato plants, demonstrating poor root development, and necrotic discolouration of the internal stem.

**Table 1.** *Fusarium oxysporum* isolates collected from processing tomato (*Solanum lycopersicum*) fields in Australia, and the GenBank accession numbers for gene sequences used in phylogenetic analyses.

| Isolate accession number<sup>a</sup> | Origin | Year | Source | GenBank Accession Number<sup>c</sup> |  |  |  |
| --- | --- | --- | --- | --- | --- | --- | --- |
|  |  |  |  | *tub2* | *cmdA* | *rpb2* | *tef1-a* |
| BRIP 5188; UOM 25087 | Stanthorpe, QLD | 1971 | n.d.<sup>b</sup> | PV746104 | PV746024 | PV746064 | PV745984 |
| BRIP 13037; UOM 25088 | Bowen, QLD | 1979 | n.d. | PV746105 | PV746025 | PV746065 | PV745985 |
| BRIP 16848; UOM 25089 | Brisbane, QLD | 1970 | n.d. | PV746106 | PV746026 | PV746066 | PV745986 |
| BRIP 17552; UOM 25090 | Stanthorpe, QLD | 1980 | n.d. | PV746107 | PV746027 | PV746067 | PV745987 |
| BRIP 76478; UOM 24033 | Himba east, VIC | 2022 | Collar | PV746108 | PV746028 | PV746068 | PV745988 |
| BRIP 76479; UOM 24034 | Himba east, VIC | 2022 | Root | PV746109 | PV746029 | PV746069 | PV745989 |
| BRIP 76480; UOM 24035 | Himba east, VIC | 2022 | Lower stem | PV746110 | PV746030 | PV746070 | PV745990 |
| BRIP 76481; UOM 24036 | Thyra, NSW | 2022 | Lower stem | PV746111 | PV746031 | PV746071 | PV745991 |
| BRIP 76482; UOM 24037 | Thyra, NSW | 2022 | Root | PV746112 | PV746032 | PV746072 | PV745992 |
| BRIP 76483; UOM 24038 | Rochester, VIC | 2022 | Collar | PV746113 | PV746033 | PV746073 | PV745993 |
| BRIP 76484; UOM 24039 | Rochester, VIC | 2022 | Collar | PV746114 | PV746034 | PV746074 | PV745994 |
| BRIP 76485; UOM 24040 | Rochester, VIC | 2022 | Collar | PV746115 | PV746035 | PV746075 | PV745995 |
| BRIP 76486; UOM 24041 | Rochester, VIC | 2022 | Collar | PV746116 | PV746036 | PV746076 | PV745996 |
| BRIP 76487; UOM 24042 | Rochester, VIC | 2022 | Root | PV746117 | PV746037 | PV746077 | PV745997 |
| BRIP 76488; UOM 24043 | Nanneella, VIC | 2023 | Collar | PV746118 | PV746038 | PV746078 | PV745998 |
| BRIP 76489; UOM 24044 | Nanneella, VIC | 2023 | Root | PV746119 | PV746039 | PV746079 | PV745999 |
| BRIP 76490; UOM 24045 | Rochester, VIC | 2023 | Collar | PV746120 | PV746040 | PV746080 | PV746000 |
| BRIP 76491; UOM 24046 | Rochester, VIC | 2023 | Root | PV746121 | PV746041 | PV746081 | PV746001 |
| BRIP 76492; UOM 24047 | Rochester, VIC | 2023 | Lower stem | PV746122 | PV746042 | PV746082 | PV746002 |
| BRIP 76493; UOM 24048 | Thyra, NSW | 2023 | Collar | PV746123 | PV746043 | PV746083 | PV746003 |
| BRIP 76494; UOM 24049 | Birganbigil, NSW | 2023 | Lower stem | PV746124 | PV746044 | PV746084 | PV746004 |
| BRIP 76495; UOM 24050 | Thyra, NSW | 2023 | Irrigation water | PV746125 | PV746045 | PV746085 | PV746005 |
| BRIP 76496; UOM 24051 | Thyra, NSW | 2023 | Irrigation water | PV746126 | PV746046 | PV746086 | PV746006 |
| BRIP 76497; UOM 24052 | Thyra, NSW | 2023 | Irrigation water | PV746127 | PV746047 | PV746087 | PV746007 |
| BRIP 76498; UOM 24053 | Thyra, NSW | 2023 | Irrigation water | PV746128 | PV746048 | PV746088 | PV746008 |
| BRIP 76499; UOM 24054 | Thyra, NSW | 2022 | Root | PV746129 | PV746049 | PV746089 | PV746009 |
| BRIP 76500; UOM 24055 | Appin, VIC | 2017 | Root | PV746130 | PV746050 | PV746090 | PV746010 |
| BRIP 76501; UOM 24056 | Burramboot, VIC | 2018 | Root | PV746131 | PV746051 | PV746091 | PV746011 |
| BRIP 76502; UOM 24057 | Leaghur, VIC | 2018 | Root | PV746132 | PV746052 | PV746092 | PV746012 |
| UOM 25002 | Burramboot, VIC | 2024 | Collar | PV746133 | PV746053 | PV746093 | PV746013 |
| UOM 25003 | Burramboot, VIC | 2024 | Root | PV746134 | PV746054 | PV746094 | PV746014 |
| UOM 25004 | Burramboot, VIC | 2024 | Collar | PV746135 | PV746055 | PV746095 | PV746015 |
| UOM 25005 | Burramboot, VIC | 2024 | Lower stem | PV746136 | PV746056 | PV746096 | PV746016 |
| UOM 25006 | Burramboot, VIC | 2024 | Root | PV746137 | PV746057 | PV746097 | PV746017 |
| UOM 25007 | Burramboot, VIC | 2024 | Collar | PV746138 | PV746058 | PV746098 | PV746018 |
| UOM 25008 | Burramboot, VIC | 2024 | Lower stem | PV746139 | PV746059 | PV746099 | PV746019 |
| UOM 25009 | Burramboot, VIC | 2024 | Collar | PV746140 | PV746060 | PV746100 | PV746020 |
| UOM 25010 | Burramboot, VIC | 2024 | Lower stem | PV746141 | PV746061 | PV746101 | PV746021 |
| UOM 25011 | Burramboot, VIC | 2024 | Root | PV746142 | PV746062 | PV746102 | PV746022 |
| UOM 25012 | Burramboot, VIC | 2024 | Collar | PV746143 | PV746063 | PV746103 | PV746023 |

<sup>a</sup> BRIP = Queensland Plant Pathology Herbarium collection. UOM = The University of Melbourne Plant Pathology collection.
<sup>b</sup> not determined
<sup>c</sup> *tub2* = beta tubulin, *cmdA* = calmodulin, *rpb2* = second largest subunit of nuclear RNA polymerase II, *tef1-α* = translation elongation factor 1-alpha.

For isolations from plant tissue, collected plants were gently washed under running tap water for approximately 5 min to remove adhering soil particles. From each plant, symptomatic sections (around 5 mm²) were excised from the roots, collar, and lower stem tissues whenever symptoms were present. In most cases, multiple tissues from the same plant were sampled to increase the likelihood of pathogen recovery, while in a few plants only one or two tissue types were available due to limited symptom development. Tissue pieces were surface-sterilised by immersion in 1% (active ingredient) sodium hypochlorite (NaClO) for 15 s, rinsed twice in sterile distilled water (30 s each), air-dried on sterile paper towels, and placed onto water agar amended with streptomycin (0.1 mg ml⁻¹; WA-S) and potato dextrose agar amended with streptomycin (0.1 mg ml⁻¹; PDA-S) (Liu *et al*., 2025). For each tissue type, three replicate plates of each medium were prepared to maximise the chance of isolating *F. oxysporum*. Plates were incubated at 24°C under continuous light and checked for fungal growth every 24 h for a week. Representative colonies displaying typical *F. oxysporum*-like morphology (white to pink mycelium and purple pigmentation) (Adhikari *et al*., 2020) from each plant organ were sub-cultured onto PDA-S.

For isolation from the irrigation water samples, each sample was dispensed into four 50 ml Falcon tubes under aseptic conditions. Although *F. oxysporum* is traditionally regarded as a soilborne pathogen, previous studies have demonstrated its capacity to survive in irrigation water and to infect host plants via water-mediated root contact (Joshi, 2018, Ullah *et al*., 2021). To recover viable pathogenic isolates from water samples, a seedling baiting approach was employed. Eight two-week-old healthy tomato seedlings grown on sterile water agar were used per water sample. Seedlings were derived from commercially imported seed certified under Australian biosecurity regulations and surface-sterilised prior to germination to minimise the risk of seed-borne contamination. The baiting protocol was adapted from Callaghan *et al*. (2022), which has been successfully applied for the recovery of plant pathogens from aquatic environments including irrigation water. Tubes were sealed with surface-sterilised parafilm, and a small opening was created to allow the seedling roots to be immersed directly in the water sample. After incubation for two weeks at room temperature, seedlings were harvested, and fungal isolations from root tissues were conducted following the same procedures described above for plant material.

Single spore isolation of each isolate was further conducted to establish pure cultures following the method described in Zhang *et al*. (2013). Four historical *F. oxysporum* isolates previously isolated from fresh tomato plants in QLD by Ramsey *et al*. (1992) were obtained from Queensland Plant Pathology Herbarium (BRIP, Brisbane) as reference Australian *F. oxysporum* isolates. Furthermore, one *F. oxysporum* isolate from 2017 and two from 2018 collected from processing tomato fields in a previous study (Callaghan, 2020) were also included (Table 1).

### 2.2. DNA extraction, PCR and sequencing

Genomic DNA from all isolates was extracted from mycelia of seven-day-old pure cultures growing on PDA. Mycelia were harvested using a sterile scalpel and transferred into 2 ml SafeLock tubes (Eppendorf, Australia), flash frozen in liquid nitrogen, and freeze-dried (Ezzi Vision, Australia) for 48-72 hours. Freeze-dried tissue was subsequently ground in a TissueLyser (Qiagen, Australia) into a fine powder using two steel balls (Sigma-Aldrich, Australia). Genomic DNA was isolated using a Qiagen DNeasy Plant Mini Kit, following the manufacturer’s instructions. The quality of the extracted DNA samples was assessed using a NanoDrop (Thermo Fisher Scientific, USA) and DNA quantification was done using a Qubit Fluorometer (Thermo Fisher Scientific). To evaluate the integrity of the DNA, agarose gel electrophoresis was conducted on 1% (wt/vol) agarose gel (in Tris Acetate EDTA buffer) stained with SYBR Safe DNA Stain (Invitrogen, USA) for visualization.

The four genomic loci, beta tubulin (*tub2*), calmodulin (*cmdA*), second largest subunit of nuclear RNA polymerase II (*rpb2*), and translation elongation factor 1-alpha (*tef1-α*), were amplified for all isolates using primer pairs T1 (5’-AACATGCGTGAGATTGTAAGT-3’)/T2 (5’-TAGTGACCCTTGGCCCAGTTG-3’) (O’Donnell & Cigelnik, 1997) for *tub2*, Cal228F (5’-GAGTTCAAGGAGGCCTTCTCCC-3’)/CAL2Rd (5’-TGRTCNGCCTCDCGGATCATCTC-3’) (Carbone & Kohn, 1999) for *cmdA*, RPB2-5F2 (5’-GGGGWGAYCAGAAGAAGGC-3’)/RPB2-7cR (5’-CCCATRGCTTGTYYRCCCAT-3’) (Liu *et al*., 1999) for *rpb2*, and EF1 (5’-ATGGGTAAGGA(A/G) GACAAGAC-3’)/EF2 (5’-GGA(G/A) GTACCAGT(G/C) ATCATGTT-3’) (O’Donnell *et al*., 1998) for *tef1-α*.

PCR reactions were adapted from the protocols described previously (Lombard *et al*., 2019, O’Donnell *et al*., 1998, Carbone & Kohn, 1999). The final concentration in each 25 μl reaction included 0.4 µM of each forward and reverse primers, 6 ng template DNA (2 ng µl^−1^), 1× PCR Master Mix (New England Biolabs, USA), and brough to volume by sterile Milli-Q water. The amplification reactions were performed in an Eppendorf thermal cycler (Eppendorf, Australia) with an initial denaturation for 9 min at 95°C; followed by 36 cycles of denaturation at 95°C for 30 s, annealing at 54°C for 45 s, and extension at 68°C for 1 minute (2 mins for *rpb2*); and a final extension at 68°C for 5 mins.

All PCR amplicons were visualised on 1% (wt vol^−1^) agarose gels, stained with SYBR Safe DNA Gel Stain, to establish amplification of a single band of expected size. PCR products were submitted to Macrogen (Seoul, South Korea) for purification and Sanger sequencing in both forward and reverse directions using the same primers used for PCR. Quality control of raw data and generation of consensus sequences were conducted in Geneious Prime 2024.0.5 (http://www.geneious.com). All consensus sequences were deposited in NCBI GenBank database (Table 1).

### 2.3. Phylogenetic analyses

Three phylogenetic analyses were conducted based on different taxonomic frameworks. These included *tef1-α* single gene analysis containing reference sequences of *F. oxysporum* from O’Donnell *et al*. (1998) and Laurence *et al*. (2012); a combined four-gene phylogeny (*tub2, cmdA, rpb2,* and *tef1-α*) with reference sequences of FOSC from Lombard *et al*. (2019); and the combined four-gene dataset with reference sequences of *F. oxysporum* from Achari *et al*. (2020).

The sequences of reference isolates were retrieved from GenBank for use in phylogenetic analyses. All sequence alignments were generated using MAFFT (Liu *et al*., 2025) plugin in Geneious Prime 2025.1.3 (https://www.geneious.com). Maximum likelihood (ML) analysis was conducted using RAxML plugin in Geneious Prime, with 1,000 pseudoreplicates and using GTRGAMMA nucleotide substitution model applied to separate partitions. Bayesian Inference (BI) analysis was conducted using MrBayes version 3.2.7a (Ronquist *et al*., 2012), and MrModeltest2 (Nylander *et al*., 2004) was used to determine the best-fit nucleotide substitutions models. Model test analysis resulted in an HKY+G model for the *tef1-α* single gene dataset containing reference sequences from O’Donnell *et al*. (1998) and Laurence et al. (2012). For the combined four-gene dataset with reference sequences from Lombard *et al*. (2019), the selected models were HKY+I+G for *tef1-α*, GTR+I+G for *rpb2*, K80+I for *cmdA*, and K80+I+G for *tub2*. For the combined four-gene dataset with reference sequences from *Achari et al.* (2020), the selected models were K80 for *tef1-α*, SYM+G for *rpb2*, K80 for *cmdA,* and HKY for *tub2*.

A neighbour-net algorithm was also used to construct a Neighbour-net network implemented in SplitsTree using the four genomic loci (*tub2, cmdA*, *rpb2,*and *tef1-α*), the uncorrected p distance and 1,000 bootstraps (Huson & Bryant, 2005). A phylogenetic network is more appropriate to depict incompatible and ambiguous signals in data sets and provide a more accurate representation of recombining data sets compared to tree-based models (Kajala *et al*., 2021). For this analyses, a subset of reference sequences were retrieved from Lombard *et al*. (2019) and Achari *et al*. (2020).

### 2.4. Sequence types

To better depict genotypic diversity within phylogenetic clades, DnaSP v. 5 was used to identify unique sequence types (haplotypes) for each of the four nuclear region (single-locus haplotypes) and for combinations of the four loci (multi-locus haplotypes) (Librado & Rozas, 2009). Estimates of the number of segregating sites (S), haplotype diversity (Hd), and nucleotide diversity (π) were conducted for each locus separately and following concatenation of loci (Librado & Rozas, 2009). Datasets as constructed were to evaluate the taxonomic position of all haplotypes and their phylogenetic relationship to recognised *F. oxysporum* species based on the four genomic loci. All sequence alignments were generated using MAFFT plugin in Geneious Prime as described before.

### 2.5. Pathogenicity bioassay

#### 2.5.1. Replicated glasshouse pathogenicity bioassay

A subset of *F. oxysporum* isolates representing different clades in the phylogenetic analyses, UOM 24033, UOM 24034, UOM24035, UOM 24036, UOM 24037, UOM 24055, UOM 24056 and UOM 24057, were selected from a total of 40 isolates for pathogenicity assays based on their four-gene haplotypes (Table S1).

Surface sterilised seeds of a commonly grown processing tomato commercial cultivar (H3402 provided by the Australian Processing Tomato Research Council) were germinated in Seed Raising Mix (Plugger111, Australian Growing Solutions Pty Ltd., Australia) for two weeks under 12 h of daylight at 25°C and 12 h of darkness at 20°C until the first pairs of true leaves emerged. For inoculation prior to planting, seedlings were dipped for 25 mins into respective conidial suspensions (10^6^ conidia mL^−1^) prepared from eight selected *F. oxysporum* isolates for inoculated treatment. Mock-inoculated control seedlings were dipped in sterile distilled water for the same duration. Inoculation procedures followed those described in Feng *et al*. (2022).

The replicated pathogenicity bioassay was conducted in the University of Melbourne Glasshouse Complex with five internal replicates for each treatment. The experiment had nine treatments (eight *F. oxysporum* isolates and a mock-inoculated control), and a total number of 45 pots. All pots contained 1.5 kg of pasteurised potting mix (Plugger 111, Australian Growing Solutions Pty Ltd.), consisting of 90% composted substances, coarse bark, and 10% sand, which were arranged in a completely randomized design. Pots were placed under a 12 h photoperiod with high-pressure sodium (HPS) light source and 25°C/20°C day/night temperatures for eight weeks. Plants were watered every other day. A complimentary Osmocote fertilizer was applied according to the recommended rate to supply essential nutrients (Osmocote, Australia). All inoculated and control (mock-inoculated) plants were checked at the end of the experiment for re-isolation of *F. oxysporum* to verify pathogenicity in accordance with Koch’s postulates (Liu *et al*., 2025). The experiment was repeated twice to confirm the replicability of the results.

#### 2.5.2. Preliminary pathogenicity assessment of an irrigation-water-derived isolate

*Fusarium oxysporum* isolate UOM 24050 recovered from irrigation water was selected for a preliminary pathogenicity assay, as all four water-derived isolates clustered within the same phylogenetic lineage and shared an identical multi-locus haplotype. The pathogenicity assay was conducted following the same glasshouse conditions described above for replicated pathogenicity experiments, using tomato cultivar H3402.

Inoculum was prepared following a modified protocol of Callaghan *et al*. (2022). For this, Trill Budgerigar millet seed mix (Trill, Australia) was soaked in tap water overnight, and subsequently autoclaved twice at 121 °C for 20 min in half-filled 250 ml plastic containers on two consecutive days. Sterile seed contents in seven containers were inoculated with *F. oxysporum* isolate UOM 24050 by transferring several agar-plugs of five or seven-day-old cultures to the plastic containers under aseptic conditions. Seven mock-inoculated containers were inoculated with clean agar plus to use for the negative control treatment. Containers were shaken thoroughly to ensure even distribution of the inoculum and incubated at room temperature for four weeks, until the millet seeds were fully colonized. Following colonization, inoculum from each treatment was air-dried in a biosafety cabinet for three days, homogenized by hand mixing, and weighed to obtain quantities for glasshouse experiment. Due to limited replication, this assay was intended to qualitatively confirm pathogenicity rather than to enable quantitative comparison.

### 2.6. Disease assessment and data analysis

Disease assessment was conducted eight weeks after inoculation on ten-week-old tomato plants. At harvest, above-ground plant height was measured and root dry weights were determined after oven-drying at 70 °C for 72 hours (Liu *et al*., 2025). Physiological parameters, including stomatal conductance (gs; mol H_2_O m^−2^ s^−1^), transpiration (E; mmol H_2_O m^−2^ s^−1^), and photosynthesis (A; μmol CO_2_ m^−2^ s^−1^), were measured using a Li-6400 XT open gas exchange system (Li-Cor Inc., Lincoln, NE, USA) (Feng *et al*., 2022). Measurements were taken from the youngest fully expanded leaves and repeated in different leaves of each plant (n = 10 per treatment, n = 90 in total). Physiological measurements were taken at weeks 2 and 6 during the glasshouse pathogenicity bioassay, due to fragile seedlings with very small leaf sizes in week 0/day 0 (the day of transplanting).

Plant root dry weight, above ground height, and physiological response data were checked for normality and homogeneity of variance. Once these assumptions were met, data were analysed using one-way ANOVA, followed by Tukey’s honestly significant difference (HSD) post hoc tests (*α* = 0.05) using Minitab 2019.1.0 (Minitab, LLC, USA) to assess significant differences between treatments.

## 3. Results

### 3.1. Sample collection and pathogen isolation

A total of 36 *F. oxysporum* isolates were obtained from the 28 tomato plants and the six irrigation water samples from processing tomato fields in in NSW and VIC, Australia (Table 1). These included thirteen isolates from collar tissues, seven from lower stem, and twelve from root, as well as four isolates from irrigation water. Together with the two isolates from 2017 and 2018 (Callaghan, 2020), these are, hereafter, referred to as the UOM isolates.

### 3.2. Phylogenetic analyses

The *tef1-α* alignment including reference isolates from O’Donnell *et al*. (1998) and *Laurence et al.* (2012) was 582 bp long and included 85 sequences. Both the ML and BI methods generated similar tree topologies; therefore, the ML tree is displayed, and BI-derived posterior probabilities were added to the ML tree (Figure 2). Most isolates clustered with reference FOSC isolates of Clade 2 and Clade 3, with low to high bootstrap support values (48 - 95%) and moderate to high Bayesian posterior probability support (0.56 - 1). Reference isolates obtained from Queensland Plant Pathology Herbarium, BRIP 5188, BRIP 13037, BRIP 16848 and BRIP 17552, and three UOM isolates, UOM 25009, UOM 24036 and UOM 24034, did not cluster with any known clades. According to the combined phylogeny, 16 isolates clustered into Clade 2 with maximum Bayesian posterior probability support, while 17 isolates clustered in Clade 3 with maximum Bayesian posterior probability (Figure 2).

**Figure 2.**
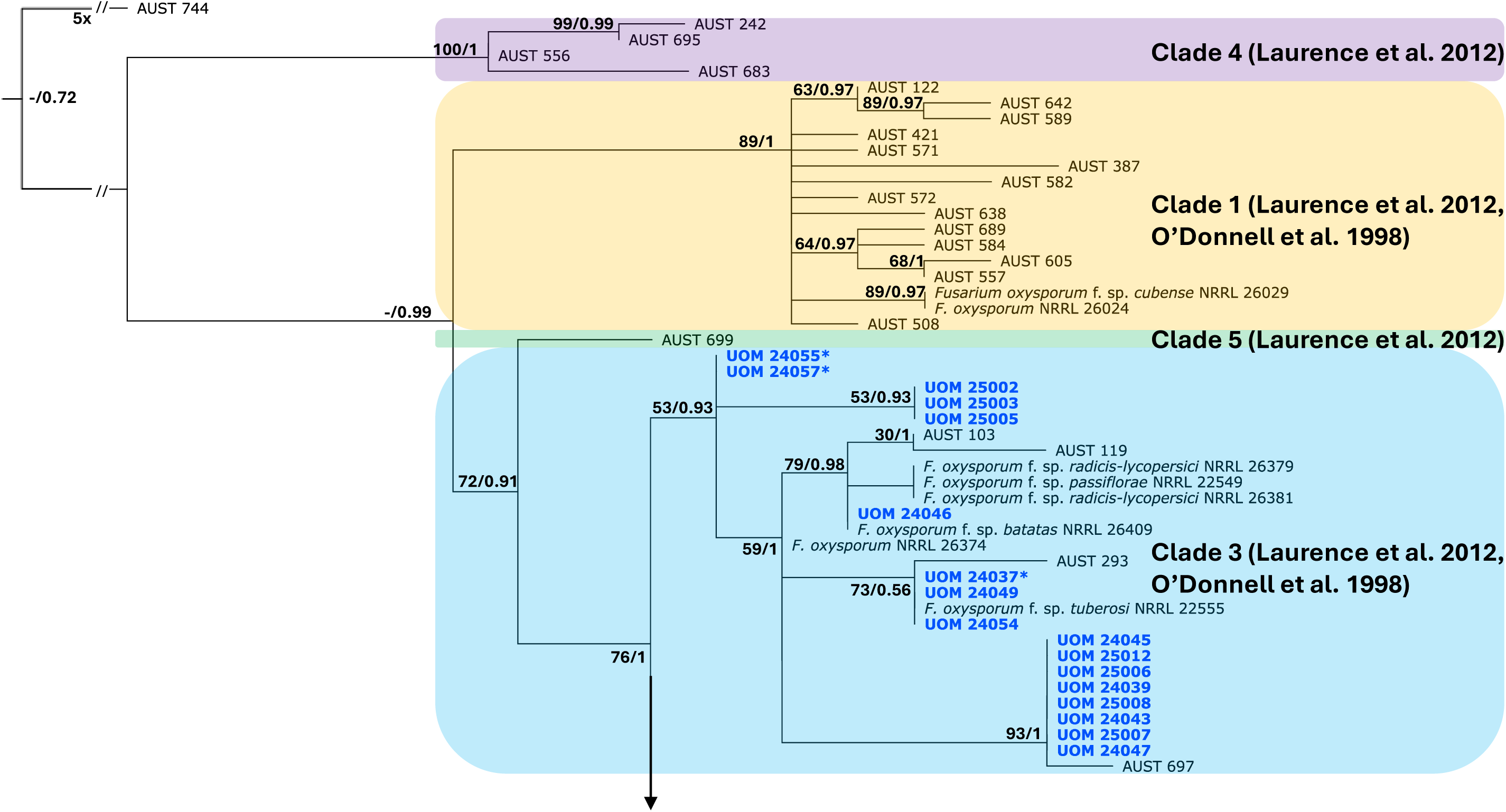
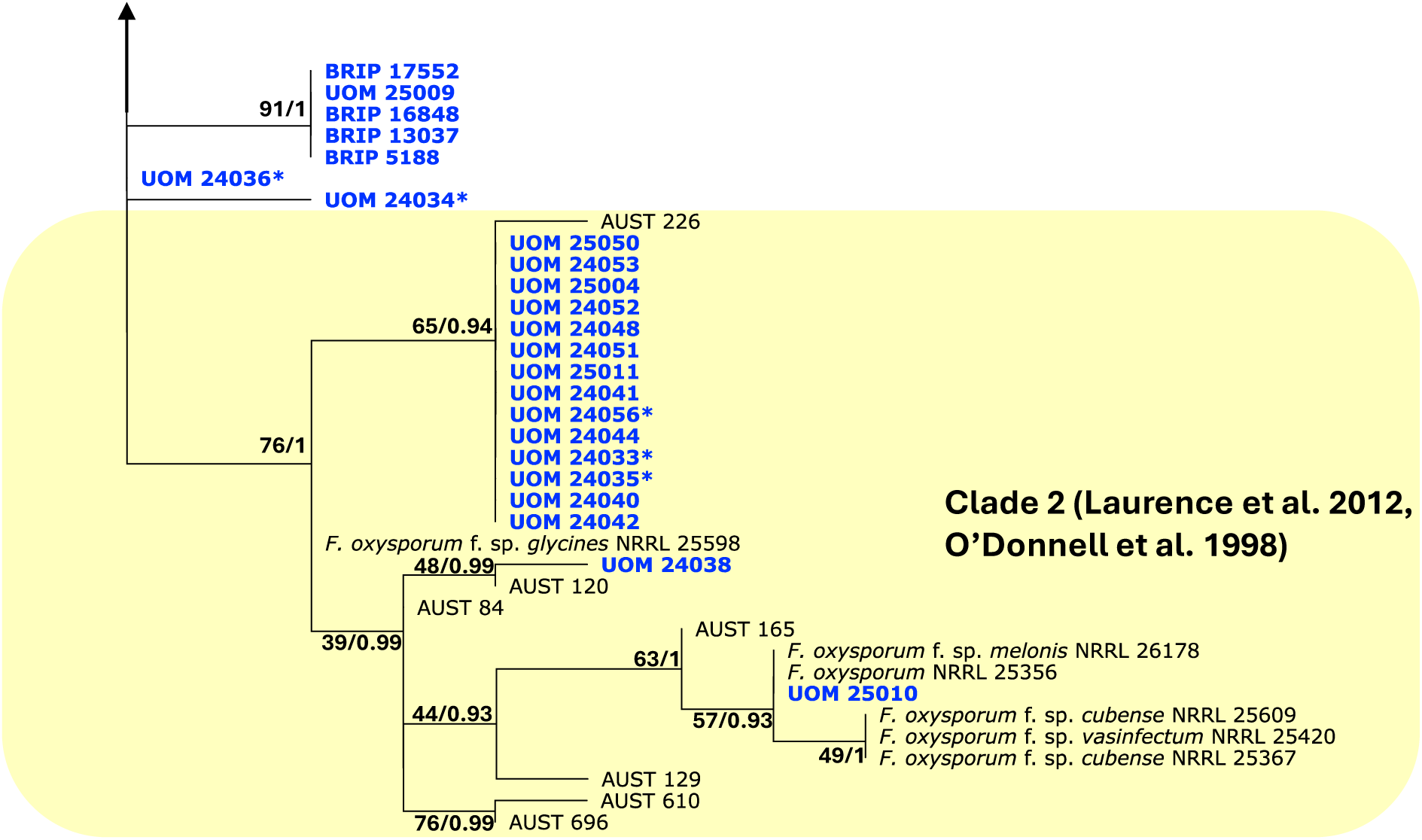
A Maximum Likelihood (ML) phylogeny of *Fusarium oxysporum* isolates inferred from the *tef1-α* alignment. The 40 isolates from tomato fields and sequenced in this study are shown in bold blue. Reference taxa were chosen to represent the species level diversity in the *F. oxysporum* species complex (FOSC) classified by O’Donnell *et al*. (1998) (NRRL isolates) and clade level diversity in the FOSC classified by Laurence *et al*. (2012) (accession numbers starting with AUST). Clade information based on Laurence *et al*. (2012) and O’Donnell *et al*. (1998) is shown on the right. The ML bootstrap support (bs) values and Bayesian posterior probability (pp) values are indicated at nodes (bs/pp). The scale bar shows the number of nucleotide changes per site. The tree was rooted with AUST 774. Isolates labelled with asterisk were used in the pathogenicity bioassay.

The four-gene dataset with reference sequences of FOSC from Lombard *et al*. (2019) included a concatenated alignment of *tub2, cmdA, rpb2,* and *tef1-α* sequences with a total length of 2,211 bp (*tub2* = 425 bp, *cmdA* = 385 bp, *rpb2* = 868 bp, and *tef1-α* = 533 bp). Given the similarity in tree structures produced by both BI and ML methods, the posterior probabilities calculated from the BI analysis were integrated into the ML tree (Figure 3). Eight major Clades were established, with the majority of UOM isolates clustering in Clades III, VI and VIII, and four UOM isolates clustering in Clade I, with no UOM isolates within Clades II, IV, V or VII. Clades III, VI and VIII, VI exhibited the highest Bayesian posterior probability support of 0.93 - 1, and Clades I, III and VIII had Bayesian support values of 0.51 - 1. All four BRIP isolates retrieved from Queensland Plant Pathology Herbarium clustered within Clade VI, with reference isolates of *F. languescens*.

**Figure 3.**
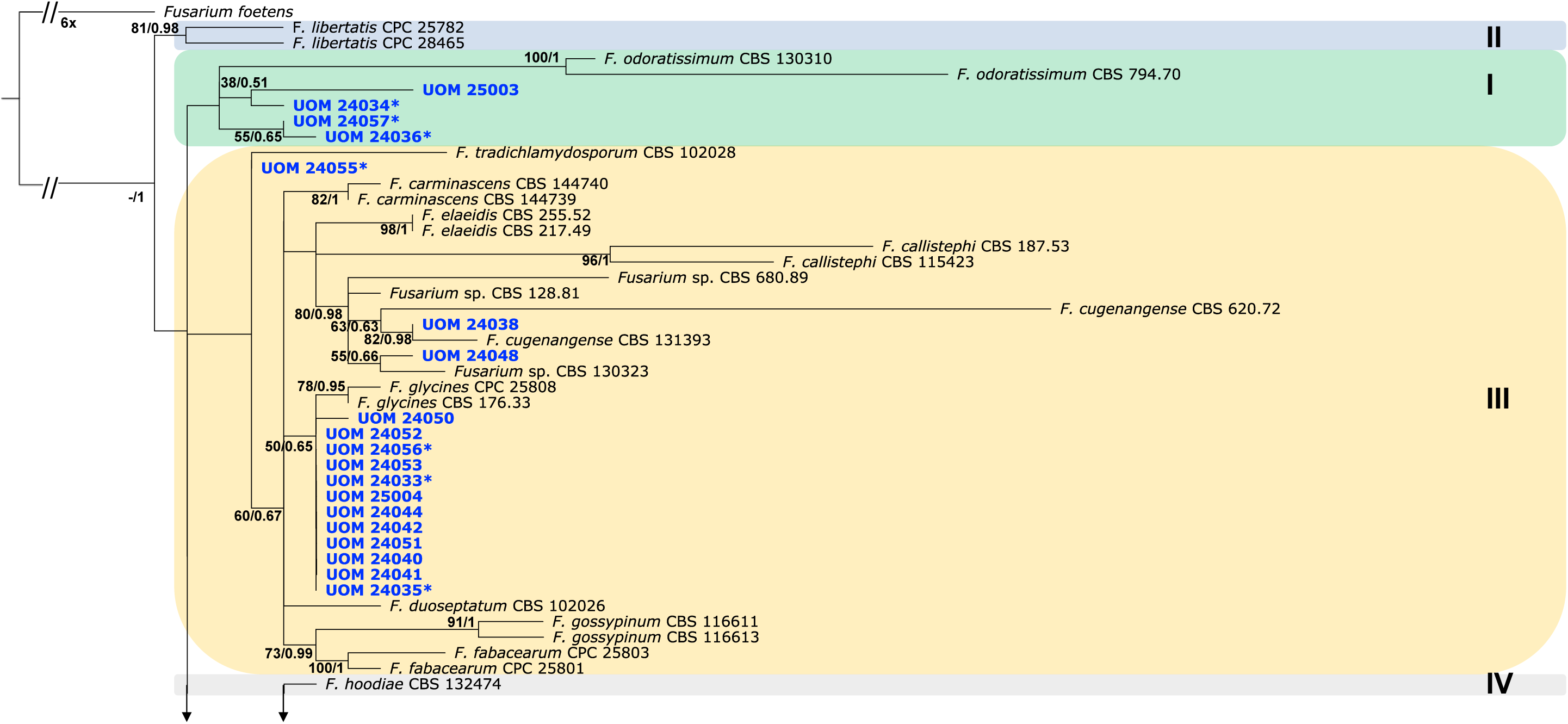
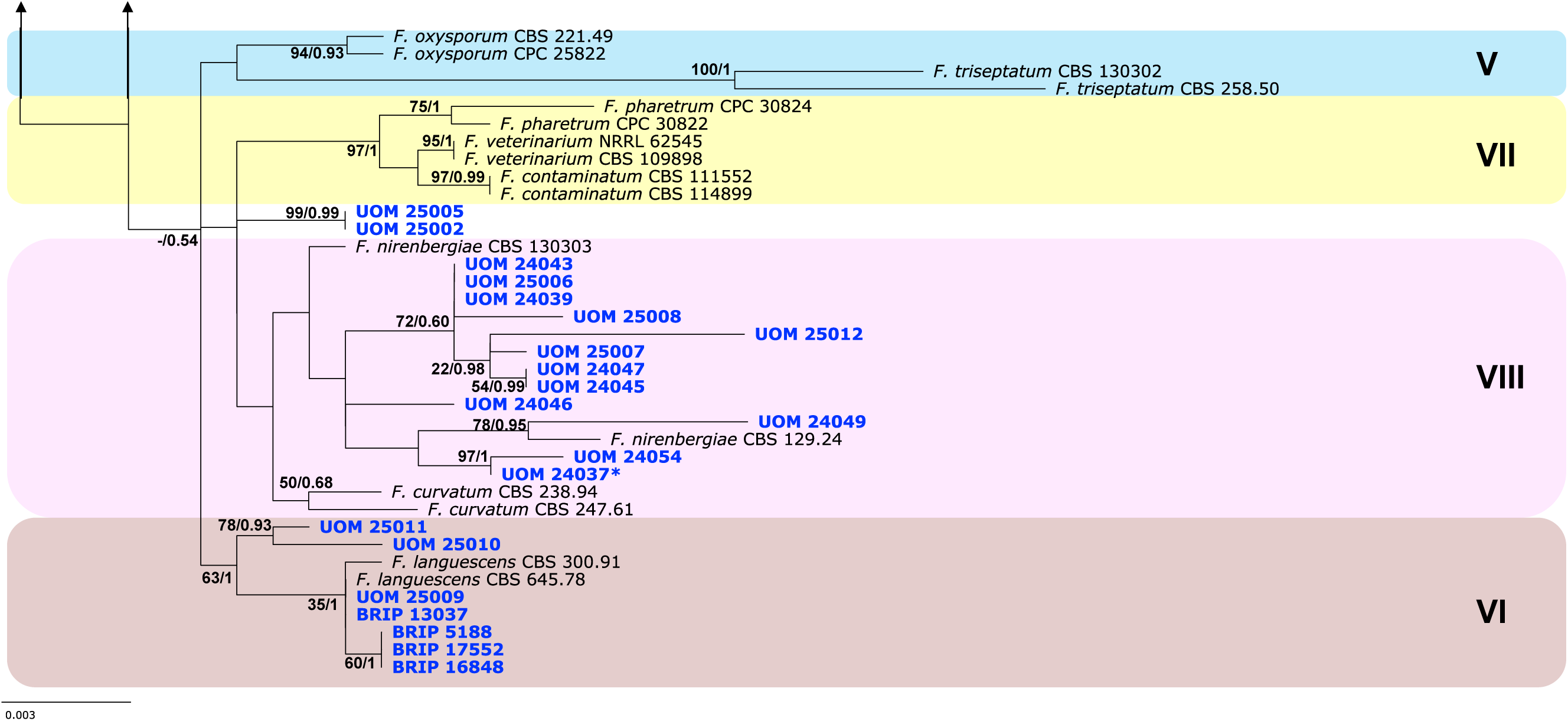
A Maximum Likelihood (ML) phylogeny of *Fusarium oxysporum* isolates inferred from concatenated alignment of *tub2, cmdA, rpb2,* and *tef1-α* sequences. Reference taxa were chosen to represent the species level diversity in the *F. oxysporum* Species Complex (FOSC) classified by Lombard *et al*. (2019). A total of 40 isolates are shown in blue bold. The ML bootstrap support values and Bayesian posterior probability values are indicated at the nodes. The scale bar shows the number of nucleotide changes per site. The tree was rooted with *Fusarium foetens*. Isolates labelled with asterisk were used in the pathogenicity bioassay

The four-gene dataset including reference sequences from Achari *et al*. (2020) resulted in a concatenated alignment of 1,959 bp (*tub2* = 413 bp, *cmdA* = 402 bp, *rpb2* = 930 bp, and *tef1-α* = 214 bp). Given the similarity in tree topologies produced by both BI and ML methods, the posterior probabilities calculated from the BI analysis were integrated into the ML tree (Figure 4). Five UOM isolates clustered in Clade 1 described by Achari *et al*. (2020) with low ML bootstrap support (63%) but maximum Bayesian posterior probability. Sixteen UOM isolates clustered into Clade 2 with ML bootstrap support of 66% and a Bayesian posterior probability of 0.99. Nineteen UOM isolates clustered in Clade 3 with high ML bootstrap support (87%) and Bayesian posterior probability value (0.83).

**Figure 4.**
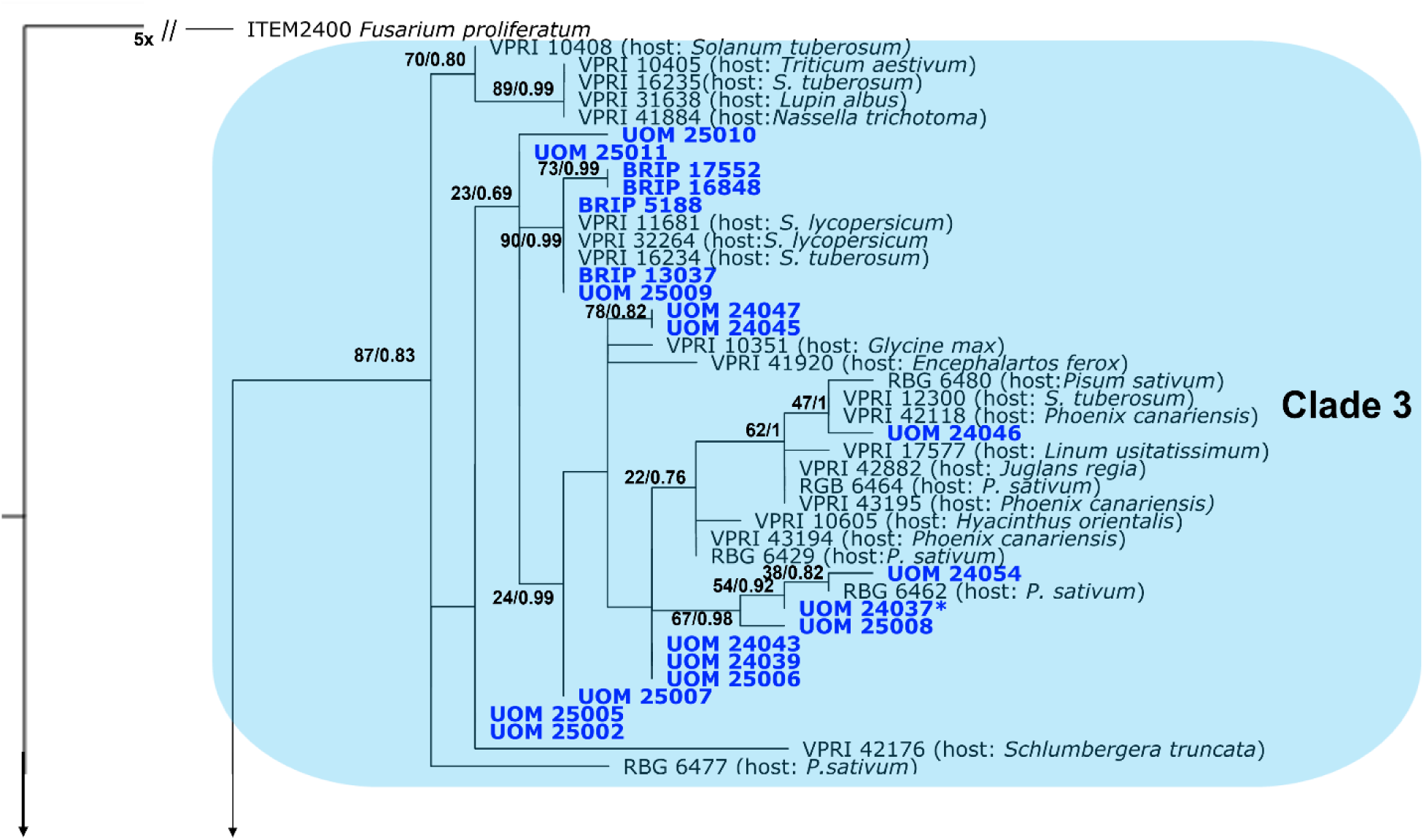
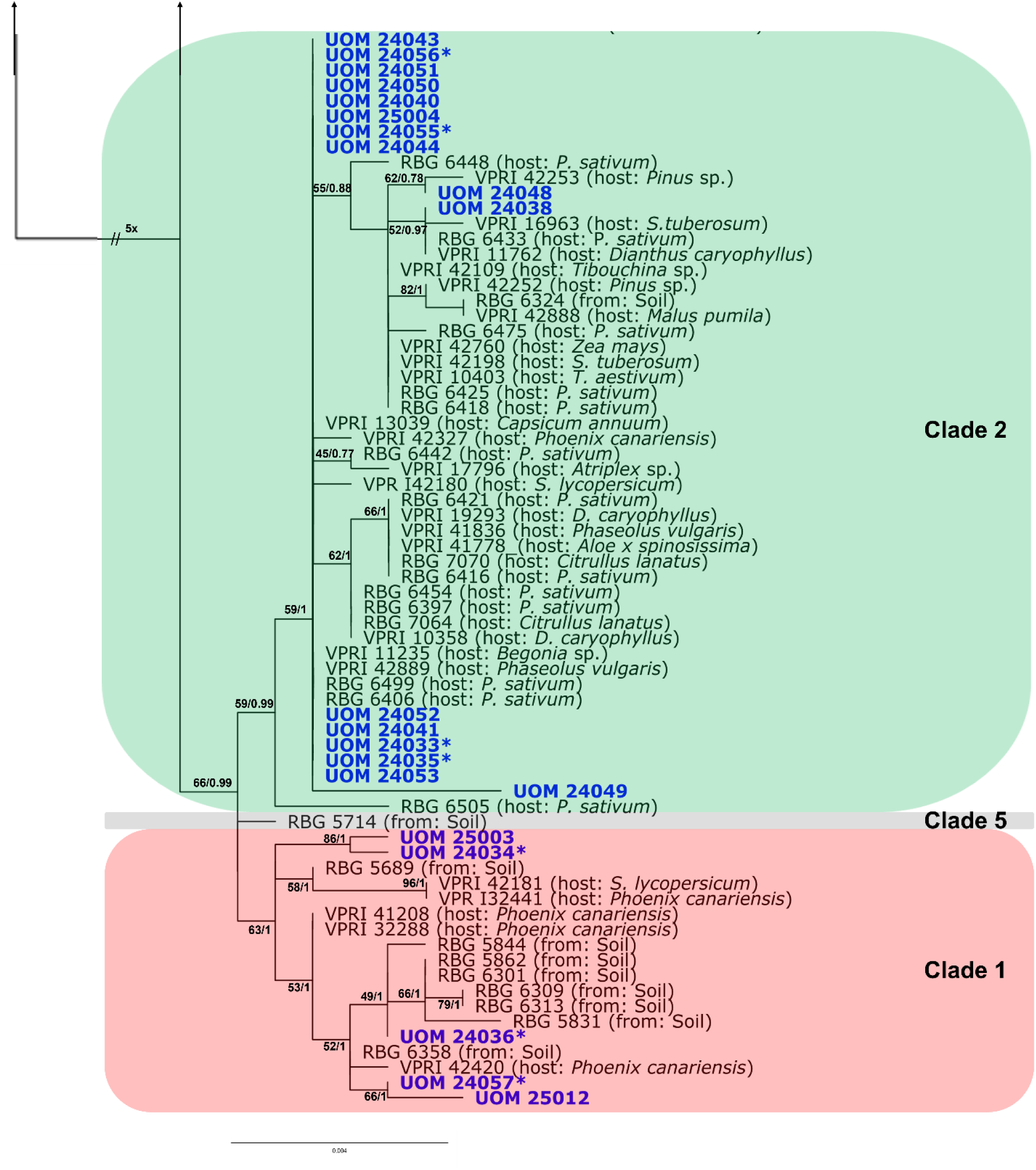
A Maximum Likelihood (ML) phylogeny of *Fusarium oxysporum* isolates inferred from the concatenated alignment of *tub2, cmdA, rpb2,* and *tef1-α*, sequences. Reference taxa were chosen to represent the species level diversity in the FOSC classified by Achari *et al*. (2020). A total of 40 isolates from tomato fields and sequenced in this study are shown in blue bold. The ML bootstrap support values and Bayesian posterior probability values are indicated at nodes. The scale bar shows the number of nucleotide changes per site. The tree was rooted with ITEM 2400 *F. proliferatum*. Isolates labelled with asterisk were used in the pathogenicity bioassay.

The Neighbour-Net phylogenetic network constructed from the four genomic loci (*tub2, cmdA*, *rpb2,* and *tef1-α*) including a subset of reference sequences from Achari *et al*. (2020) and Lombard *et al*. (2019) further resolved the FOSC isolates into three well-defined clades (Figure 5). Clade 1 (red) was composed of the tomato-associated reference isolate VPRI 41281 *Solanum lycopersicum*, and five UOM isolates, with moderate support value of 56%. Clade 2 (green) included *F. eleaeidis*, *F. fabacearum*, and *F. carminascens*, which contained 16 UOM isolates, supported by bootstrap values ranging from 56% to 74%. Clade 3 (blue) included *F. pharertum*, *F. veterinarum*, and *F. curvatum*, along with numerous UOM and all BRIP reference isolates, with strong support for major internal nodes (87%, 95%, and 97%). Lower support values were observed at deeper nodes, reflecting reticulation among clades.

**Figure 5.**
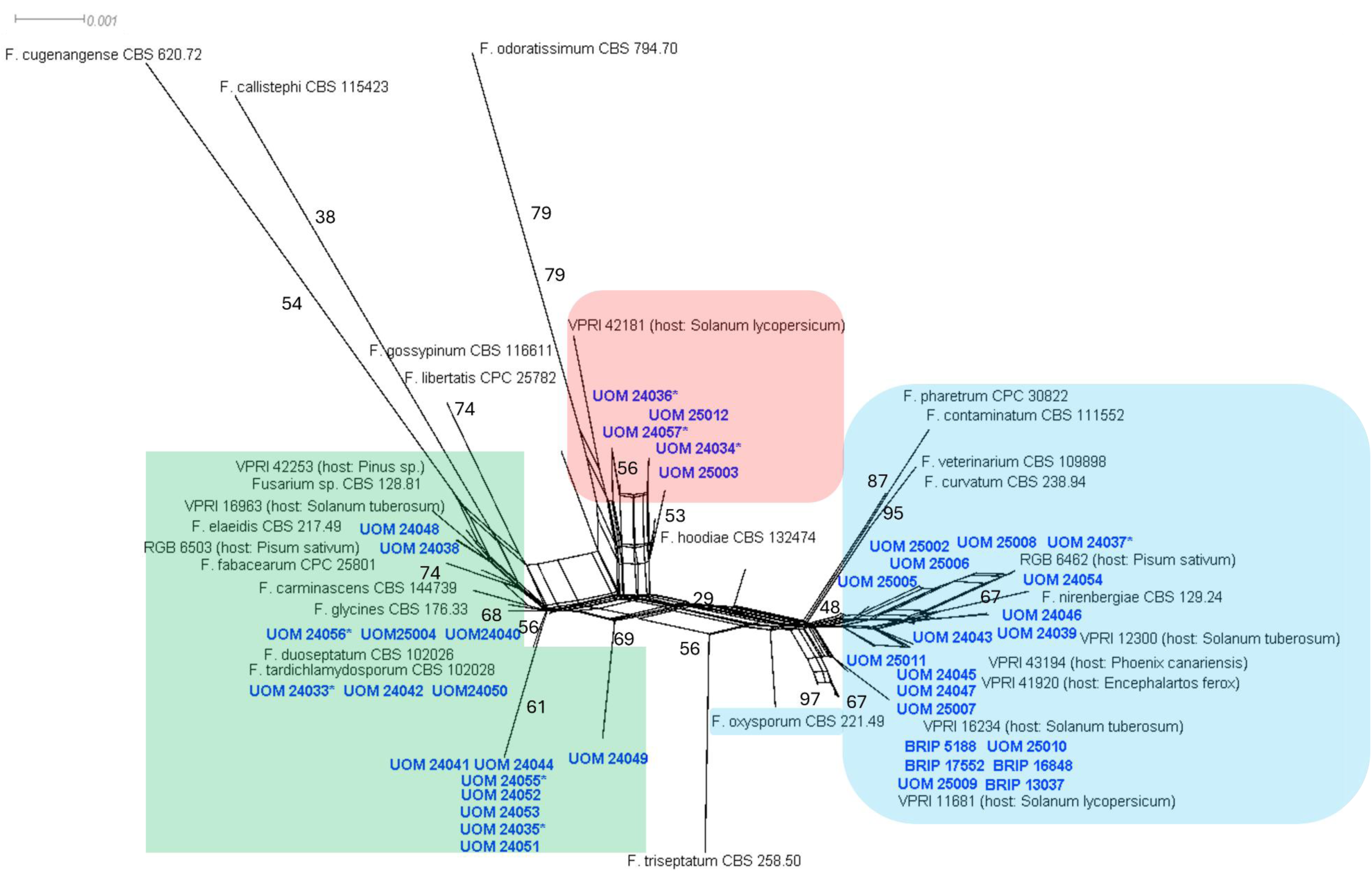
Phylogenetic relationships among *Fusarium oxysporum* isolates based on concatenation of four genomic loci (*tub2, cmdA, rpb2,* and *tef1-α*). Shown in blue are isolates collected from processing tomato fields in NSW and VIC, and the four reference isolates (BRIP) from fresh tomato in Qld. Shown in black are reference isolates from Lombard *et al*. (2019) and Achari *et al*. (2020). The scale bar represents the expected changes per site. Isolates with asterisk were used in the pathogenicity bioassay. Red, green, and blue boxes represent Clades 1, 2, and 3 based on the phylogenetic analyses proposed by Achari *et al*. (2020).

### 3.3. Multi-locus haplotypes and estimates of genetic variation

Sequencing the *tub2, cmdA*, *rpb2,* and *tef1-α*, regions for 36 UOM *F. oxysporum* isolates collected from processing tomato fields in NSW and VIC, as well as the four reference BRIP isolates, detected 56 polymorphic sites and 27 haplotypes in the combined data of the four gene sequences and a total alignment length of 2,200 bp (Tables 2 and S1). For each gene sequence, the most frequent haplotype for *tub2* was *tub2*-Hap 1 that included 28 isolates with all BRIP isolates present, followed by *tub2*-Hap 2 that contained 5 isolates. For *cmdA*, the most frequent haplotype was *cmdA*-Hap 1 with 20 isolates in total, followed by *cmdA*-Hap 2 which included 17 isolates. For *rpb2,* the most frequent haplotype was *rpb2*-Hap 3 that contained 14 isolates, followed by *rpb2*-Hap 6, which included eight isolates. The most frequent *tef1-α* haplotype (*tef1-α-*Hap 5) included 14 isolates, followed by *tef1-α*-Hap 2 that contained eight isolates. The highest degree of sequence polymorphism was detected in the *tef1-α* region with a nucleotide diversity of 0.009 (Table 2). Detailed haplotypes of the four genomic loci of the 40 isolates are provided in Supporting information Table S1.

**Table 2.** Summary statistics of DNA polymorphism in four nuclear loci (*tub2, cmdA, rpb2* and *tef1-α*) sequenced for 40 *F. oxysporum* isolates from tomato sequenced in this study.

| Locus | *tub2* | *cmdA* | *rpb2* | *tef1-a* | combined |
| --- | --- | --- | --- | --- | --- |
| **No. of sites** | 415 | 362 | 893 | 530 | 2,200 |
| <sup>a</sup>**S** | 3 | 3 | 20 | 20 | 46 |
| <sup>b</sup>**H** | 3 | 4 | 11 | 10 | 24 |
| <sup>c</sup>**H<sub>d</sub>** | 0.304 | 0.581 | 0.832 | 0.818 | 0.908 |
| <sup>d</sup>**π** | 0.0009 | 0.0018 | 0.0056 | 0.0091 | 0.0049 |
| **D<sup>e</sup>** | -1.0624 n.s. | -0.1432 n.s. | 0.2349 n.s. | -0.1234 n.s. | -0.0769 n.s. |

<sup>a</sup> S = Number of polymorphic sites (number of parsimony informative sites).
<sup>b</sup> H = number of haplotypes.
<sup>c</sup> H<sub>d</sub> = haplotype diversity.
<sup>d</sup> π = Nucleotide diversity defined as the average number of nucleotide differences per site between any two randomly chosen DNA sequences.
<sup>e</sup> D = Tajima's D; n.s. = not significant (p > 0.05)

### 3.4. Pathogenicity and aggressiveness of *F. oxysporum* on processing tomato

Koch’s postulates were satisfied. All inoculated processing tomato plants exhibited reduced growth compared with the control, whereas control plants remained healthy. The pathogen was consistently re-isolated from inoculated plants and confirmed as *F. oxysporum* based on culture morphology identical to the original isolates. Above ground height of inoculated plants ranged from 35.60 ± 0.35 to 42.5 ± 0.21 cm, compared with 47.9 ± 0.39 cm in the control. Root development was similarly reduced, with dry root weights ranging from 1.23 ± 0.19 to 3.89 ± 0.11 g in inoculated plants and 7.01 ± 0.28 g in the control (Table 3). These reductions were significant (p < 0.05) regardless of the genetic background of the isolates. Furthermore, plant physiological responses were significantly reduced in all inoculated plants compared with the control (mock-inoculated treatment) at both measurement times (weeks 2 and 6; p < 0.05; Table 4). Photosynthesis ranged from 9.85 ± 0.16 to 13.39 ± 0.20 μmol CO₂ m⁻² s⁻¹ at week 2 and 3.79 ± 0.13 to 12.67 ± 0.13 μmol CO₂ m⁻² s⁻¹ at week 6 (control: 14.47 ± 0.29 and 14.94 ± 0.12, respectively). Transpiration ranged from 3.29 ± 0.10 to 4.78 ± 0.11 mmol H₂O m⁻² s⁻¹ at week 2 and 2.00 ± 0.09 to 4.53 ± 0.11 mmol H₂O m⁻² s⁻¹ at week 6 (control: 5.07 ± 0.14 and 6.15 ± 0.10), while stomatal conductance ranged from 0.43 ± 0.11 to 0.90 ± 0.07 mol H₂O m⁻² s⁻¹ at week 2 and 0.31 ± 0.01 to 0.67 ± 0.01 mol H₂O m⁻² s⁻¹ at week 6 (control: 1.25 ± 0.11and 0.98 ± 0.04). Similar trends were observed at both time points, with all reductions being statistically significant. Among the eight *F. oxysporum* isolates tested in the pathogenicity bioassay, UOM 24034 showed the highest root dry weight, above ground height, and physiological parameters among inoculated plants, indicating it was the least aggressive isolate. In contrast, UOM 24055 showed the lowest values for these traits, making it the most aggressive isolate of those tested (Tables 3 and 4). Additionally, the representative *F. oxysporum* isolate UOM 24050 recovered from irrigation water was pathogenic on tomato. Infected seedlings developed symptoms consistent with *F. oxysporum* infection, including leaf chlorosis, defoliation, and early wilting compared with mock-inoculated controls. Because this assay was conducted with limited replication, data were not included for quantitative comparison due to differing experimental conditions.

**Table 3.** Mean (± standard error) values of root dry weight and above-ground height of processing tomato plants eight weeks after inoculation with *Fusarium oxysporum* isolates. Different letters indicate significant differences among means (α = 0.05).

| Treatment | Isolate haplotype <sup>a</sup> | Isolate source | Root dry weight (g) ± standard error (SE) | Above ground height (cm) <sup>b</sup> ± SE |
| --- | --- | --- | --- | --- |
| Control | - | - | 7.0118<sup>A</sup> ± 0.2750 | 47.90<sup>A</sup> ± 0.3912 |
| UOM 24033 | 16 | Collar | 1.5704<sup>C</sup> ± 0.0878 | 37.80<sup>CDE</sup> ± 0.3614 |
| UOM 24034 | 24 | Root | 3.8850<sup>B</sup> ± 0.1050 | 42.50<sup>B</sup> ± 0.2131 |
| UOM 24035 | 4 | Lower stem | 1.9240<sup>C</sup> ± 0.0522 | 36.80<sup>DE</sup> ± 0.3451 |
| UOM 24036 | 5 | Lower stem | 1.8614<sup>C</sup> ± 0.0538 | 39.80<sup>BCD</sup> ± 0.2652 |
| UOM 24037 | 18 | Root | 2.1412<sup>C</sup> ± 0.1242 | 39.40<sup>BCD</sup> ± 0.1691 |
| UOM 24055 | 2 | Root | 1.2270<sup>C</sup> ± 0.1918 | 35.60<sup>E</sup> ± 0.2909 |
| UOM 24056 | 16 | Root | 1.7126<sup>C</sup> ± 0.1839 | 37.20<sup>DE</sup> ± 0.2648 |
| UOM 24057 | 1 | Root | 1.6266<sup>C</sup> ± 0.1985 | 40.60<sup>BC</sup> ± 0.2843 |

<sup>a</sup> Haplotypes refers to the unique sequence types for the four-locus haplotype for each isolate (Tables 2 and S1)
<sup>b</sup> Different letters A - E show significant differences between treatments (rows) based on ANOVA and the Tukey honestly significant difference (HSD) post hoc tests ( $\alpha = 0.05$ ; $p < 0.05$ )

**Table 4.** Mean (± standard error) values of tomato plant physiological parameters in response to inoculation with *Fusarium oxysporum* isolates at two- and six-weeks post-inoculation.

| Treatment | Photosynthesis ($\mu\text{mol CO}_2 \text{ m}^{-2} \text{ s}^{-1}$) ± standard error (SE) | | Stomatal Conductance ($\text{mol H}_2\text{O m}^{-2} \text{ s}^{-1}$) ± SE | | Transpiration ($\text{mmol H}_2\text{O m}^{-2} \text{ s}^{-1}$) ± SE | |
| --- | --- | --- | --- | --- | --- | --- |
|  | W2 | W6 | W2 | W6 | W2 | W6 |
| Control | 14.47<sup>A</sup> ± 0.2923 | 14.94<sup>A</sup> ± 0.1205 | 1.25<sup>A</sup> ± 0.1103 | 0.98<sup>A</sup> ± 0.0420 | 5.07<sup>A</sup> ± 0.1368 | 6.15<sup>A</sup> ± 0.0951 |
| UOM 24033 | 9.89<sup>F</sup> ± 0.1551 | 5.48<sup>F</sup> ± 0.1511 | 0.50<sup>CD</sup> ± 0.0570 | 0.41<sup>E</sup> ± 0.0095 | 3.65<sup>D</sup> ± 0.0617 | 3.21<sup>E</sup> ± 0.0810 |
| UOM 24034 | 13.39<sup>B</sup> ± 0.1962 | 12.67<sup>B</sup> ± 0.1336 | 0.90<sup>B</sup> ± 0.0700 | 0.67<sup>B</sup> ± 0.0119 | 4.55<sup>BC</sup> ± 0.1555 | 4.53<sup>B</sup> ± 0.1107 |
| UOM 24035 | 10.71<sup>DEF</sup> ± 0.1800 | 10.18<sup>D</sup> ± 0.1916 | 0.59<sup>CD</sup> ± 0.0136 | 0.57<sup>C</sup> ± 0.0098 | 4.42<sup>BC</sup> ± 0.1290 | 3.90<sup>CD</sup> ± 0.0370 |
| UOM 24036 | 11.04<sup>D</sup> ± 0.0519 | 9.93<sup>D</sup> ± 0.1390 | 0.57<sup>CD</sup> ± 0.0144 | 0.56<sup>CD</sup> ± 0.0109 | 4.13<sup>C</sup> ± 0.0522 | 3.89<sup>CD</sup> ± 0.0450 |
| UOM 24037 | 12.24<sup>C</sup> ± 0.1990 | 11.46<sup>C</sup> ± 0.1437 | 0.70<sup>BC</sup> ± 0.0237 | 0.65<sup>B</sup> ± 0.0053 | 4.78<sup>AB</sup> ± 0.1093 | 4.22<sup>BC</sup> ± 0.0620 |
| UOM 24055 | 9.85<sup>F</sup> ± 0.1559 | 3.79<sup>G</sup> ± 0.1280 | 0.48<sup>CD</sup> ± 0.0103 | 0.31<sup>F</sup> ± 0.0057 | 3.29<sup>D</sup> ± 0.0962 | 2.00<sup>F</sup> ± 0.0906 |
| UOM 24056 | 10.81<sup>DE</sup> ± 0.1837 | 10.00<sup>D</sup> ± 0.2770 | 0.60<sup>CD</sup> ± 0.0113 | 0.57<sup>C</sup> ± 0.0111 | 4.29<sup>C</sup> ± 0.0332 | 3.98<sup>C</sup> ± 0.0563 |
| UOM 24057 | 9.99<sup>EF</sup> ± 0.2675 | 9.10<sup>E</sup> ± 0.0490 | 0.43<sup>D</sup> ± 0.1093 | 0.50<sup>D</sup> ± 0.0064 | 4.26<sup>C</sup> ± 0.0689 | 3.60<sup>D</sup> ± 0.0820 |

Different letters A - F show significant differences between treatments (rows) based on ANOVA and the Tukey honestly significant difference (HSD) post hoc tests ( $\alpha = 0.05$ ; $p < 0.05$ )

Despite the variability in aggressiveness among isolates, only two inoculated plants exhibited minor internal lesions in the lower stems under glasshouse conditions (Figure 6). No other visible lesions or discolorations were observed. Nevertheless, all inoculated plants showed significant reductions in above ground growth, root production, and physiological parameters, demonstrating that the tested isolates were pathogenic under glasshouse conditions.

**Figure 6.**
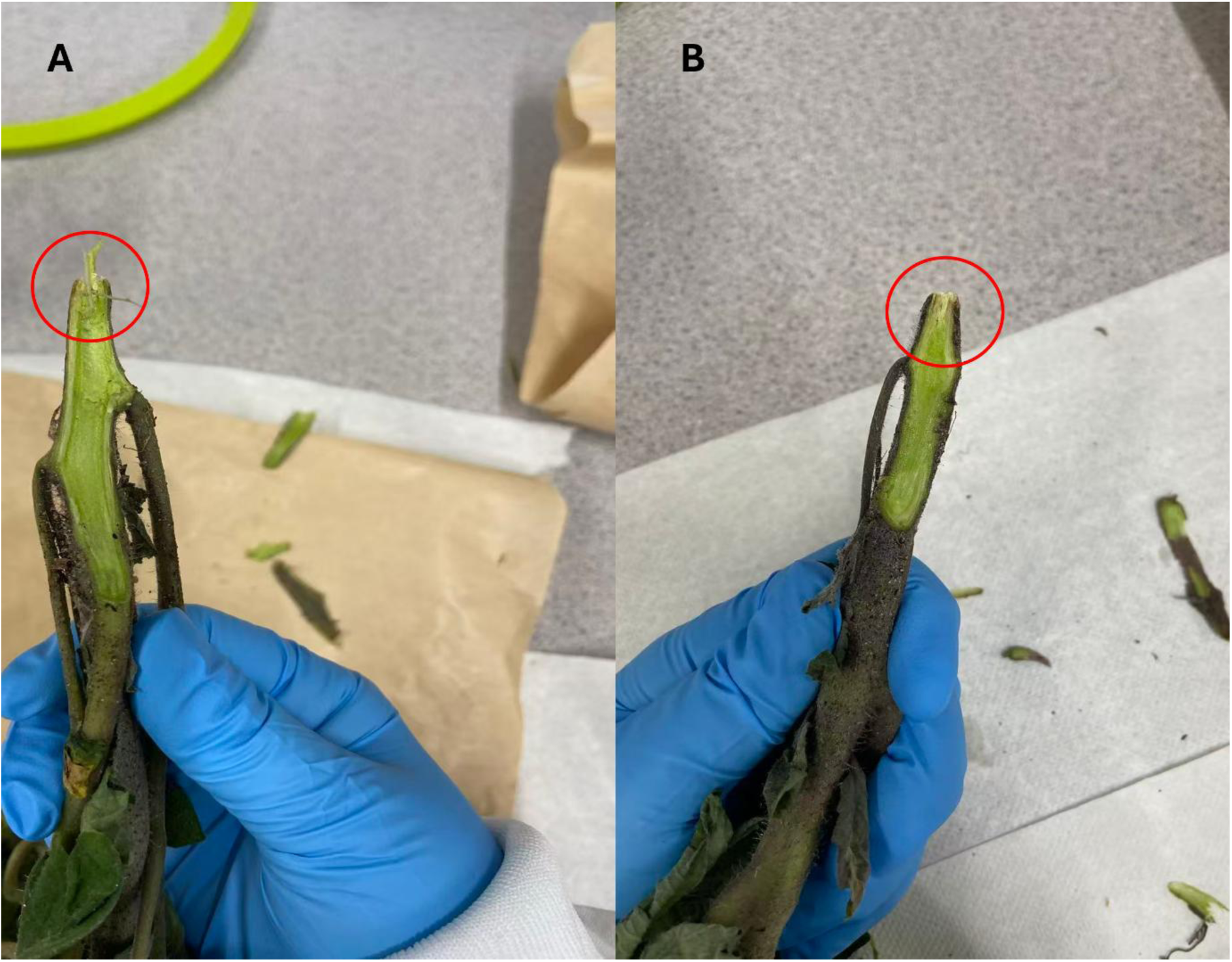
Minor internal lesions in the lower stems of two inoculated tomato plants at the end of the replicated glasshouse bioassay. (A) Plant inoculated with UOM 24056, (B) plant inoculated with UOM 24055.

## 4. Discussion

Phylogenetic analyses of four genomic loci in *Fusarium oxysporum* Species Complex (FOSC) isolates collected from processing tomato fields in Australia revealed substantial genetic diversity and genetic exchange among clades, highlighting the complexity of their taxonomic placement. Isolates were placed across different clades within FOSC, suggesting the potential involvement of multiple phylogenetic species in yield decline in processing tomato.

Species assignment within the FOSC is complicated by the complex evolutionary history of isolates as revealed through extensive reticulation, likely driven by frequent horizontal gene transfer and recombination, and also the use of differing taxonomic frameworks in FOSC taxonomy. Based on the four-gene phylogeny proposed by Lombard *et al*. (2019), the isolates were assigned to at least eight species within the FOSC. In contrast, the systems proposed by O’Donnell *et al*. (1998) and Laurence *et al*. (2012) placed these isolates within two of the three (O’Donnell *et al*., 1998) and five clades (Laurence *et al.,* 2012), respectively, while the application of the methodology proposed by Achari *et al*. (2020) assigned them to four clades. Despite variations among different taxonomic frameworks, multi-locus phylogenetic and coalescent-based analyses including nuclear, mitochondrial, and genome-wide datasets, have consistently aligned with earlier studies that support the division of FOSC into three major clades (Aoki *et al*., 2014, Brankovics *et al*., 2017, Laurence *et al*., 2012), two of which encompassed the tomato-associated isolates.

In addition to the genetic diversity observed through tree-based analyses, the phylogenetic network analysis further highlighted the substantial genetic complexity within the FOSC with three well-defined clades. These results supported that the tested isolates represent diverse phylogenetic lineages within the FOSC and align more closely with the three-clade classification than with the eight-clade taxonomy proposed by Lombard *et al*. (2019). The extensive reticulation among isolates indicates exchange of genetic material among clades potentially through recombination and horizontal gene transfer. This is consistent with previous studies, which have reported similar reticulated network patterns reflecting recombination and gene flow within the FOSC (Hoogendoorn *et al*., 2018, Fayyaz *et al*., 2023). The detection of genetic exchange across diverse clades underscores the evolutionary flexibility of *F. oxysporum* and poses further challenges for species delimitation, diagnostics, and disease management strategies. While our results show the presence of at least two phylogenetic species within the tomato-associated FOSC isolates, we refrain from describing separate species or *formae speciales* due to the absence of evidence for ecological or host separation among the isolates and the lack of consistency across different approaches.

Multi-locus haplotype analysis revealed that all four isolates obtained from irrigation water samples, UOM 24050, UOM 24051, UOM 24052, and UOM 24053, were assigned to Hap-16, the most frequently observed haplotype isolated from tomato plants based on the combined four-gene sequence dataset (Table S1). The occurrence of a dominant haplotype in irrigation water indicates that recycled irrigation water from a single processing tomato field may serve as a source of *F. oxysporum* inoculum contributing to disease epidemics. Although *F. oxysporum* is traditionally regarded as a soil-borne pathogen, members of the FOSC are capable of persisting in aquatic environments (Graça *et al*., 2016), potentially through the formation of stress-tolerant propagules such as chlamydospores, association with organic debris, or survival on plant residues present in irrigation systems (Palmero *et al*., 2011). This suggests that contaminated irrigation water may contribute to the dissemination of clonal lineages within and across fields. Such a route of spread aligns with previous studies that have identified irrigation water as a critical factor in the epidemiology of *F. oxysporum* (Ullah *et al*., 2021), further underscoring the potential role of water-mediated dispersal of the dominant haplotype around processing tomato growing regions.

The genetic diversity across the four gene regions analysed varied considerably, with *rpb2* and *tef-1α* showing greater diversity than *cmdA* and *tub2*. Specifically, *rpb2* exhibited the highest number of polymorphic sites (S), highest haplotype diversity (Hd), and second highest nucleotide diversity (π). *tef-1α* followed with the second highest Hd and π values, indicating a similarly high level of genetic variability. In contrast, *cmdA* and *tub2* displayed lower S, Hd, and π values, pointing to lower resolution power of these loci for within-species diversity analyses. Previous studies have also found that *tef-1α* exhibited high sequence type diversity whereas *rpb2* provided high-resolution phylogenetic information, indicating significant genetic diversity (Nephalela-Mavhunga *et al*., 2021).

The replicated pathogenicity bioassay conducted under controlled glasshouse conditions confirmed that all eight isolates were pathogenic to the commercial processing tomato cultivar tested in Australia. Infected plants exhibited significantly reduced dry matter accumulation and impaired physiological responses. Reductions in photosynthesis rate, stomatal conductance, and transpiration were observed in inoculated plants and are consistent with the growth reduction recorded in this study. These physiological changes are commonly associated with vascular wilt diseases, where xylem colonisation and impairment of water transport lead to reduced gas exchange (Deans *et al*., 2020), stomatal closure (Muir, 2020), and diminished carbon assimilation (Verma *et al*., 2009). In the present study, declines in physiological performance broadly aligned with reductions in above ground biomass and root development, supporting their use as indicators of disease impact rather than as standalone diagnostic traits.

Although pathogenicity testing of irrigation-water-derived *F. oxysporum* isolates was limited in scope and replication, qualitative assessment of a representative Hap-16 isolate UOM 24050 confirmed its ability to cause disease on tomato. This finding supports the epidemiological relevance of irrigation systems as potential sources of viable inoculum associated with the dominant haplotype identified in processing tomato fields. Because this assessment was conducted under different experimental conditions and without accompanying physiological measurements, the results were not intended for quantitative comparison with replicated pathogenicity assays, but rather to demonstrate pathogenic potential of water-associated isolates.

Notably, the commercial cultivar tested here (cv. H3402), although reported to exhibit resistance to Fusarium wilt in the United States (Stewart, 2023), did not show resistance to the Australian *F. oxysporum* isolates. Therefore, further screening and evaluation of imported processing tomato cultivars using Australian isolates are essential to identify and develop effective resistance against the diverse Australian *F. oxysporum* populations.

In addition, various degrees of aggressiveness were observed among the eight tested isolates, irrespective of their phylogenetic placement or genetic variation observed among the isolates across the three phylogenetic frameworks. This highlights that phylogenetic classification, which reflects genetic relatedness based on housekeeping loci, does not necessarily predict pathogenicity or aggressiveness. Pathogenicity depends on phenotypic assays, rather than sequence similarity alone. Consequently, assessing disease potential cannot rely solely on genomic sequences, and experimental evaluation of host–pathogen interactions remains essential. These findings are consistent with previous studies showing that genetic divergence among FOSC isolates did not always correspond to phenotypic or pathogenic differentiation sufficient to justify species-level distinctions (Manawasinghe *et al*., 2021).

Last but not least, classical race classifications of *Fusarium oxysporum* f. sp. *lycopersici* have been instrumental for resistance breeding; however, these frameworks are largely based on a limited number of well-characterised Northern Hemisphere lineages (Swett *et al*., 2023, Srinivas *et al*., 2019). Increasing evidence suggests that tomato-pathogenic *F. oxysporum* populations can be phylogenetically diverse and may fall outside canonical FOL clades while retaining high virulence. In such cases, race-associated markers and effector profiles may not reliably reflect evolutionary relationships or pathogenic potential. These observations reinforce the need for phylogeny-based approaches to understand pathogen diversity, particularly in under-studied regions such as Australia.

In conclusion, *Fusarium oxysporum* Species Complex was demonstrated to be a dominant pathogen affecting processing tomato crops in Victoria and New South Wales, Australia. The inconsistent phylogenetic groupings derived from three analytical frameworks using identical genomic loci support the polyphyletic nature of the pathogen. A replicated glasshouse pathogenicity bioassay confirmed the field-observed growth reductions, although characteristic disease symptoms such as wilting, root rot, and collar lesions were not consistently reproduced. Notably, all FOSC isolates tested caused statistically similar growth suppression on the most commonly used commercial cultivar H3402, indicating a lack of complete resistance regardless of isolate phylogeny. The successful isolation of *F. oxysporum* from irrigation water also highlights its potential role as a possible inoculum source in field epidemiology. These findings underscore the need for further investigation into the evolutionary origins and population structure of *F. oxysporum* Species Complex obtained in Australian processing tomato fields, along with its epidemiological dynamics, to improve host resistance selection and inform more effective, regionally tailored disease management strategies.

## Acknowledgement

The authors thank the Australian Processing Tomato Research Council (APTRC) for funding this project, we appreciate the tremendous help and support from Matthew Stewart and Ann Morrison from Australian Processing Tomato Industry. The first author gratefully appreciates the support of Melbourne Research Scholarship provided by the University of Melbourne.

## Data Availability Statement

Data available on request from the authors.

