## Supplementary Table S1 for "Genetic characterisation and aggressiveness of the *Fusarium oxysporum* Species Complex in tomato plants and irrigation water from Australian processing fields"

**Supporting information**

**Table S1.** Details of the 40 *Fusarium oxysporum* isolates collected from processing tomato fields in NSW and VIC, and reference isolates obtained from Queensland Plant Pathology Herbarium (BRIP). Beta tubulin (*tub2*), calmodulin (*cmdA*), the second largest subunit of nuclear RNA polymerase II (*rpb2*), and translation elongation factor 1-alpha (*tef1-α*), were generated for all isolates. The unique sequence types (haplotypes) for each sequenced region and the four-locus haplotype for each isolate are shown.

| **Isolate** | **Single locus haplotypes** | | | | **Four-locus haplotypes** |
|---|---|---|---|---|---|
| | ***tub2*** | ***cmdA*** | ***rpb2*** | ***tef1-α*** | |
| BRIP 5188 | Haplotype 1 | Haplotype 1 | Haplotype 6 | Haplotype 9 | Haplotype 23 |
| BRIP 13037 | Haplotype 1 | Haplotype 2 | Haplotype 6 | Haplotype 9 | Haplotype 22 |
| BRIP 16848 | Haplotype 1 | Haplotype 1 | Haplotype 6 | Haplotype 9 | Haplotype 23 |
| BRIP 17552 | Haplotype 1 | Haplotype 1 | Haplotype 6 | Haplotype 9 | Haplotype 23 |
| UOM 24033 | Haplotype 1 | Haplotype 1 | Haplotype 3 | Haplotype 5 | Haplotype 16 |
| UOM 24034 | Haplotype 1 | Haplotype 1 | Haplotype 2 | Haplotype 8 | Haplotype 21 |
| UOM 24035 | Haplotype 1 | Haplotype 1 | Haplotype 3 | Haplotype 5 | Haplotype 16 |
| UOM 24036 | Haplotype 1 | Haplotype 3 | Haplotype 1 | Haplotype 10 | Haplotype 24 |
| UOM 24037 | Haplotype 1 | Haplotype 2 | Haplotype 10 | Haplotype 1 | Haplotype 4 |
| UOM 24054 | Haplotype 2 | Haplotype 2 | Haplotype 10 | Haplotype 1 | Haplotype 5 |
| UOM 24038 | Haplotype 3 | Haplotype 1 | Haplotype 5 | Haplotype 6 | Haplotype 18 |
| UOM 24039 | Haplotype 2 | Haplotype 2 | Haplotype 8 | Haplotype 2 | Haplotype 10 |
| UOM 24040 | Haplotype 1 | Haplotype 1 | Haplotype 3 | Haplotype 5 | Haplotype 16 |
| UOM 24041 | Haplotype 1 | Haplotype 1 | Haplotype 3 | Haplotype 5 | Haplotype 16 |
| UOM 24042 | Haplotype 1 | Haplotype 1 | Haplotype 3 | Haplotype 5 | Haplotype 16 |
| UOM 24043 | Haplotype 2 | Haplotype 2 | Haplotype 8 | Haplotype 2 | Haplotype 10 |
| UOM 24044 | Haplotype 1 | Haplotype 1 | Haplotype 3 | Haplotype 5 | Haplotype 16 |
| UOM 24045 | Haplotype 1 | Haplotype 2 | Haplotype 9 | Haplotype 2 | Haplotype 12 |

| | | | | | |
|---|---|---|---|---|---|
| UOM 24046 | Haplotype 2 | Haplotype 4 | Haplotype 11 | Haplotype 4 | Haplotype 15 |
| UOM 24047 | Haplotype 2 | Haplotype 2 | Haplotype 9 | Haplotype 2 | Haplotype 9 |
| UOM 24048 | Haplotype 1 | Haplotype 1 | Haplotype 4 | Haplotype 5 | Haplotype 20 |
| UOM 24049 | Haplotype 2 | Haplotype 4 | Haplotype 3 | Haplotype 1 | Haplotype 3 |
| UOM 24050 | Haplotype 1 | Haplotype 1 | Haplotype 3 | Haplotype 5 | Haplotype 16 |
| UOM 24051 | Haplotype 1 | Haplotype 1 | Haplotype 3 | Haplotype 5 | Haplotype 16 |
| UOM 24052 | Haplotype 1 | Haplotype 1 | Haplotype 3 | Haplotype 5 | Haplotype 16 |
| UOM 24053 | Haplotype 1 | Haplotype 1 | Haplotype 3 | Haplotype 5 | Haplotype 16 |
| UOM 24055 | Haplotype 1 | Haplotype 1 | Haplotype 3 | Haplotype 1 | Haplotype 2 |
| UOM 24056 | Haplotype 1 | Haplotype 1 | Haplotype 3 | Haplotype 5 | Haplotype 16 |
| UOM 24057 | Haplotype 1 | Haplotype 1 | Haplotype 1 | Haplotype 1 | Haplotype 1 |
| UOM 25002 | Haplotype 1 | Haplotype 2 | Haplotype 7 | Haplotype 3 | Haplotype 14 |
| UOM 25003 | Haplotype 1 | Haplotype 2 | Haplotype 2 | Haplotype 3 | Haplotype 13 |
| UOM 25004 | Haplotype 1 | Haplotype 1 | Haplotype 3 | Haplotype 5 | Haplotype 16 |
| UOM 25005 | Haplotype 1 | Haplotype 2 | Haplotype 7 | Haplotype 3 | Haplotype 14 |
| UOM 25006 | Haplotype 1 | Haplotype 2 | Haplotype 8 | Haplotype 2 | Haplotype 8 |
| UOM 25007 | Haplotype 1 | Haplotype 2 | Haplotype 6 | Haplotype 2 | Haplotype 11 |
| UOM 25008 | Haplotype 1 | Haplotype 2 | Haplotype 10 | Haplotype 2 | Haplotype 7 |
| UOM 25009 | Haplotype 1 | Haplotype 2 | Haplotype 6 | Haplotype 9 | Haplotype 22 |
| UOM 25010 | Haplotype 1 | Haplotype 2 | Haplotype 6 | Haplotype 7 | Haplotype 19 |
| UOM 25011 | Haplotype 1 | Haplotype 2 | Haplotype 6 | Haplotype 5 | Haplotype 17 |
| UOM 25012 | Haplotype 1 | Haplotype 2 | Haplotype 1 | Haplotype 2 | Haplotype 6 |
